# Growth balance analysis models of cyanobacteria for understanding resource allocation strategies

**DOI:** 10.1101/2023.04.18.537335

**Authors:** Sajjad Ghaffarinasab, Martin J. Lercher, Hugo Dourado

## Abstract

Cyanobacteria have emerged as attractive microbial cell factories, as they can convert atmospheric CO_2_ and sunlight into valuable chemicals. To increase their growth and productivity, one should aim to optimize the allocation of limited cellular resources across different metabolic processes. Here, we developed two growth balance analysis (GBA) models for the cyanobacterium *Synechocystis* sp. PCC 6803. In its biological assumptions, the models closely related to an existing coarse-grained model, while its mathematical formulation is heavily streamlined. We show that the GBA models provide virtually identical predictions about cellular resource allocation among photosynthesis, carbon metabolism, and the protein translation machinery under different environmental conditions as the previous, mathematically more involved model. Our model also captures the effects of photodamage on proteome allocation and the resulting growth rates. We further show how the GBA model can be easily extended to include more reactions, leading to a second GBA model capable of new predictions about the cellular resource allocation. Balanced growth models of the type presented here can easily expanded to include more biological details, providing a useful toolbox for the understanding of the physiological capabilities of cyanobacteria, their allocation of cellular resources, and the potential of their bioengineering for optimized biomass production.

## Introduction

Photosynthetic cyanobacteria are the only prokaryotes which are capable of oxygenic photosynthesis, converting CO_2_ and sunlight to biomass. Cyanobacteria exhibit higher photosynthetic efficiency as well as faster growth and are more accessible to genetic manipulations than plants and microalgae. These features make them an important model organism for designing microbial cell factories(Santos-Merino, Yun, & Ducat, 2023). While a vast amount of experimental high-throughput data is available - such as genomics, transcriptomics and proteomics (Babele, Kumar, & Chaturvedi, 2019; Jahn et al., 2018; Matthias et al., 2014; Zavřel et al., 2019)-a mechanistic understanding of cyanobacterial resource allocation from simple principles remains an ongoing challenge in biotechnology.

Computational models allow the study of different phenotypes, growth conditions and biological principles that are not easily accessible to experiments. Recently, various linear approaches in computational biology have been used to study resource allocation of organisms, for example genome-scale Models of Metabolism and Macromolecular Expression, ME-models(O’Brien, Lerman, Chang, Hyduke, & Palsson, 2013), Resource Balance Analysis, RBA (Goelzer, Fromion, & Scorletti, 2011), and genome-scale models with enzymatic constraints using kinetic and omics data, GECKO (Sánchez et al., 2017). These approaches consider the production cost of macromolecules for catalyzing each reaction by estimating the kinetic rate laws as a linear relationship between fluxes and the concentration of their catalysts, ignoring metabolite concentrations and how they influence fluxes via the saturation of the catalysts(Dourado & Lercher, 2020).

Alternative modeling approaches have been developed to account for non-linear kinetic rate laws, thus also including metabolite concentrations explicitly in the models.(Molenaar, Van Berlo, De Ridder, & Teusink, 2009) initially established a small-scale heterotrophic cell model, where the allocation of its resources follows directly from the optimization of its growth rate under basic physiological constraints. (Burnap, 2015) and (Faizi, Zavř003Bel, Loureiro, Červený, & Steuer, 2018) later extended this modeling approach to simulate photoautotrophic resource allocation, including specific properties and physiology, such as photodamage and carbon-cycling. Nonlinear optimization problems are however difficult to be solved numerically for large scale models (Wortel, Noor, Ferris, Bruggeman, & Liebermeister, 2018), and nonlinear kinetics have been currently accounted for only in small, simplified cell models (Burnap, 2015; Faizi et al., 2018; Jahn et al., 2018; Molenaar et al., 2009). Recent studies have shown how the mathematical formulation of such nonlinear problems can be greatly simplified, potentially facilitating the simulation of larger nonlinear cell models(Dourado & Lercher, 2020; Dourado, Liebermeister, Ebenhöh, & Lercher, 2022).

In this work, we develop as a proof of concept two Growth Balance Analysis (GBA) models: the first, based on an existing mechanistic model of the cyanobacterium *Synechocystis* sp. PCC 6803 by (Faizi et al., 2018), and a second extended model with more reactions detailing the photosynthetic apparatus. We show how the GBA approach reduces the mathematical modeling to a minimum set of assumptions, while retaining the same fundamental prediction capabilities of the original model by (Faizi et al., 2018). We then show how the second GBA model is easily built from the first model without changes in the mathematical formalism, resulting in new predictions that are consistent with existing experimental data for *Synechocystis* sp. PCC 6803.

## RESULTS

### First model: a simple cyanobacterium model including photoinhibition

Our first GBA model of cyanobacterium *Synechocystis* sp. PCC 6803 (hereafter *Synechocystis*) is based on the recent resource allocation model of (Faizi et al., 2018)(hereafter also referred simply as “Faizi model”), including four reactions representing main cellular functions: carbon transport (T), metabolism and carbon assimilation (M), ribosome and protein translation (R), and photosynthetic unit (P). The main purpose of such models is to reduce all the cellular complexity to only few components and reactions that still provide important insights into the main patterns of cellular behavior. This simplification is done by combining multiple essential enzymes and metabolic pathways into single catalytic units. Coarse-grained models are helpful tools to identify metabolic tradeoffs under various circumstances without requiring much information about the organisms(Molenaar et al., 2009).

(Faizi et al., 2018) models photosynthesis as the conversion between an “inactive” protein unit *P*^0^ and an “active” protein unit *P*^∗^, absorbing light from the environment and producing energy “e” as a result. The simple GBA framework does not account for proteins as substrates of reactions, thus our model treats photosynthesis as a simple transport reaction “PSU” importing energy “e” into the system. We also simplify the Faizi model by ignoring carbon passive diffusion through the cell membrane, and only account for an active import of carbon by the protein transporter “T”. A GBA model is defined by the triple (M,τ,ρ), where **M** denotes a mass fraction matrix (i.e. an internal stoichiometric matrix normalized by corresponding molecular masses, including a column “R” for a ribosome reaction producing proteins, and a row “a” for the total protein concentration), τ denotes reaction turnover times determined by some given kinetic rate laws, and ρ denotes biomass density. For any given GBA model, a simple set of equations determining the balanced growth problem can be derived from first principles (Dourado et al., 2022). These equations depend only on the mass balance of reactions contained in the matrix **M**, kinetic parameters, and cell density data, which are only a subset of all parameters used in the *Synechocystis* model from (Faizi et al., 2018). The optimal cellular state at an environment defined by the external concentration of carbon and light intensity *x* = (*x*_*C*_, *x*_*L*_) is then found as the one maximizing the balanced growth rate μ under the constraints of mass conservation, reaction kinetics and fixed cell density. The details of our first GBA model is presented on **Table 1**, and **Figure 1** presents the corresponding schematic.

**Table 1.** The parameters, properties, and equations defining our first GBA model for *Synechocystis*.

|  |  |
| --- | --- |
| Model parameters | $\mathbf{M}$: mass fraction matrix<br>$k_{cat}^j$: turnover number of reaction $j$ [$\text{h}^{-1}$]<br>$K_m^j$: Michaelis constant of metabolite $m$ in reaction $j$ [$\text{g L}^{-1}$]<br>$I_n^j$: Inhibition constant for external concentration $n$ in reaction $j$ [$\text{g L}^{-1}$]<br>$\rho$: cellular density [$\text{g L}^{-1}$] |
| System properties | $v_j$: flux of reaction $j$ [$\text{g L}^{-1} \text{h}^{-1}$]<br>$\mu$: growth rate [$\text{h}^{-1}$]<br>$\mathbf{c}$: reactant concentration vector (including concentrations $c^m$ of metabolites and the total protein concentration $c^a$) [$\text{g L}^{-1}$]<br>$p_j$: protein concentration of $j$ [$\text{g L}^{-1}$]<br>$\tau_j$: turnover time of $j$ [$\text{h}$] |
| Mass conservation constraint | $\mathbf{M}\mathbf{v} = \mu\mathbf{c}$ ,<br>where |
| | $\mathbf{M} = \begin{bmatrix} 1 & 0 & -0.7 & 0 \\ -0.2 & 1 & -0.3 & -0.1 \\ 0 & 0 & 1 & -0.9 \\ 0 & 0 & 0 & 1 \end{bmatrix}$ |
| Reaction kinetics | $p_j = v_j \tau_j(\mathbf{c}, \mathbf{x})$ |
| constraint | where |
| | $\tau_j(\mathbf{c}, \mathbf{x}) = \frac{1}{k_{cat}^j} \prod_m \left( \frac{c_m}{K_m^j + c_m} \right)^{-1} \prod_n \left( \frac{x_n}{K_n^j + x_n} \right)^{-1} \left( \frac{I_n^j}{I_n^j + x_n} \right)^{-1}$ |
| Total protein constraint | $c_a = \sum_j p_j$ |
| Cellular density | $\rho = c_a + \sum_m c^m$ |
| constraint |  |

**Figure 1.**
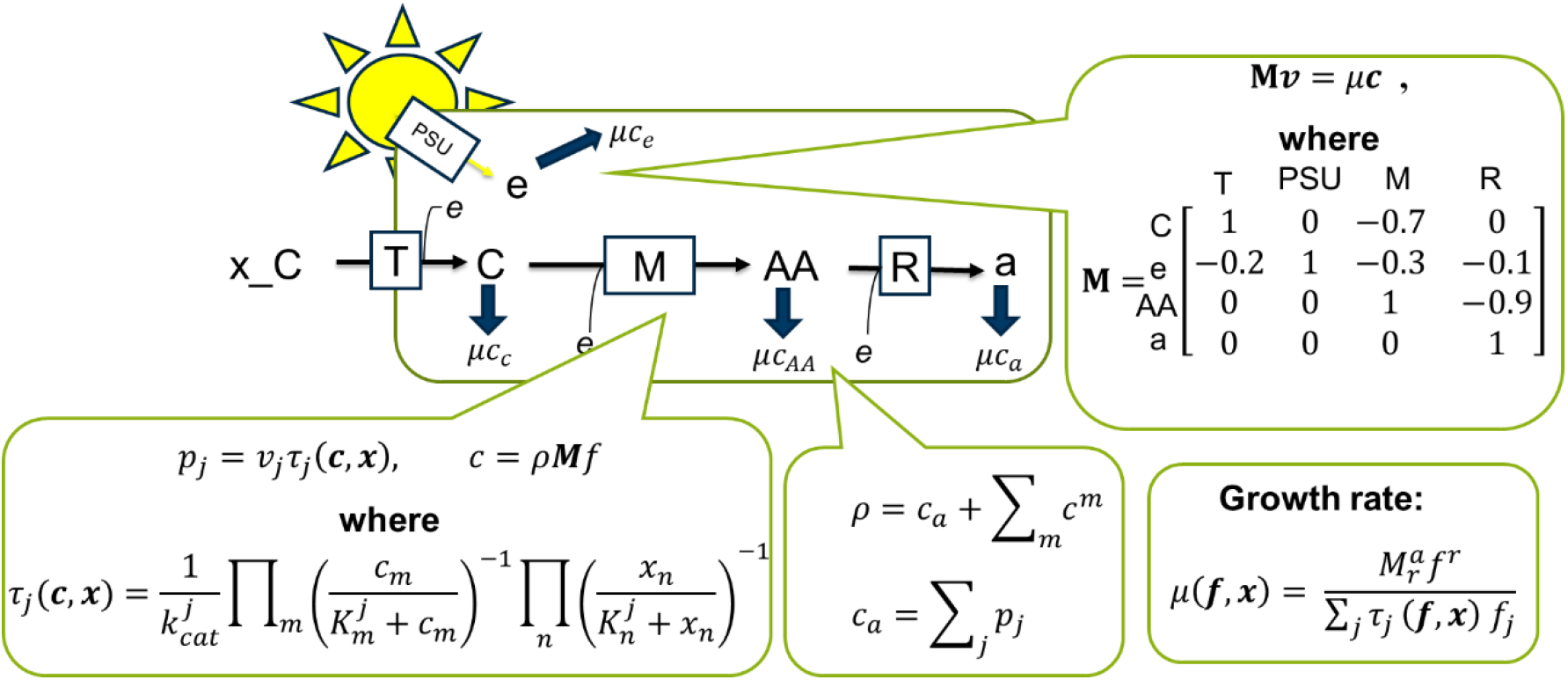
Overview of the first growth balance analysis (GBA) model for *Synechocystis*. External “metabolites” (light and external inorganic carbon) are imported by the transporters PSU, T, so a metabolic reaction “M” converts the internal C,e into aminoacids “AA”, which is used by the ribosome “R” to produce all protein “a” in the model. The protein “a” is assumed to be instantly distributed into the four reactions (PSU, T, M, R) so their protein catalyst maintains its concentration constant at balanced growth. Mass conservation of internal reactants (C, e, AA) at balanced growth is enforced by the equation **M***v* = μ***c***, relating fluxes *v*, growth rate μ, and internal mass concentrations ***c*** = (*c*_*C*_, *c*_*e*_, *c*_AA_, *c*_*a*_), accounting for the dilution by growth. Each reaction *j* is catalyzed by a specific protein with concentration *p*_*j*_ and a turnover rate τ_*j*_(***c, x***), which is determined by kinetic rate laws and depends on the internal concentrations ***c*** and external concentrations *x* = (*x*_*C*_, *x*_*e*_) of reactants involved in the reaction. The fixed cellular density ρ constrains the sum of all metabolite concentrations *c*_m_ and total protein concentration *c*_*a*_ (itself defined as the sum of all individual protein concentrations *p*_*j*_). The growth rate optimization problem at given environment ***x*** = (*x*_*C*_, *x*_*I*_) can be entirely formulated on flux fractions ***f*** = *v*/μρ, greatly simplifying analytical and numerical studies(Dourado et al., 2022).

We model the photoinhibition of growth by accounting for an inhibition term in a general kinetic rate law (Liebermeister & Klipp, 2006). We consider that this inhibition affects only the ribosome reaction, to account for how the photodamage of proteins degraded into amino acids demand an increase in the protein production.

In Faizi model, the optimization problem is also defined by maximizing the growth rate at external environmental conditions *x* = (*x*_*C*_, ***x***_*L*_) of carbon concentration and light intensity, under the constraints of mass conservation, reaction kinetics, fixed total concentration of components (Faizi et al., 2018). The rate laws are assumed to be simple irreversible Michaelis-Menten kinetics, except for a phenomenological equation for the photosynthesis reaction. The resulting nonlinear optimization problem is defined on the variables β_*j*_ defined as the proportion of ribosomes allocated to translate each protein *j*. The equations also depend on the internal concentrations of carbon, energy, and amino acids *c*_*C*_, *c*_*e*_, *c*_AA_, and their first derivatives in time; the resulting set of 7 ODEs is treated as a separated numerical problem, which is first solved before the numeric optimization of growth rate by the variables ***β***.

In the GBA framework, the entire optimization problem is completely defined by two algebraic equations in terms of *flux fractions **f*** : = *v*/ μ ρ; the objective function is given by the growth rate function μ (*f, x*),

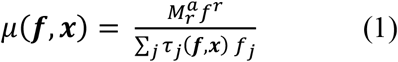

where 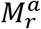 is the entry in the matrix **M** corresponding to the ribosome column “r” and total protein row “a”, and the constraint on cellular density

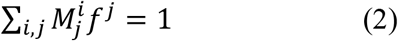

where we have sums for all rows *i* and all columns *j* of **M**. Equation (1) already encodes all the constraints in Table 1 (see Methods and (Dourado et al., 2022) for details), except for the density constraint captured by equation (2) in terms of ***f***. The vector ***f*** determines all system properties listed on Table 1, including the fluxes *v*, the concentrations *c* (including metabolite concentrations

*c*_m_ and total protein concentration *c*_*a*_)

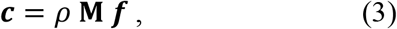

and the protein concentrations *p*^*j*^ allocated to each reaction *j*

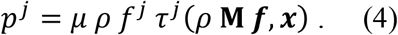

The reaction turnover times τ are defined by rate laws (here irreversible Michaelis-Menten with inhibition) depending on metabolite mass concentrations *c*_m_, which are in turn uniquely determined by equation (3). For transport reactions at the cell surface, their turnover times τ also depend on external concentrations (*x*) determining the environment condition.

We note that Faizi model employs molar units, while in GBA all the equations are mass normalized and thus simplified. In our study, we utilized molecular weights of metabolites to normalize the mass fraction matrix M, as well as the Michaelis constants 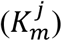 and turnover numbers 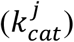 of the enzymes. Specifically, each 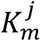 values in [mol L^-1^] were multiplied by the corresponding molecular mass in [g mol^-1^], resulting in [g L^-1^]. Similarly, the 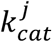 values [h^-1^] were multiplied by the product molecular mass and divided by the respective enzyme molecular mass, resulting in [h^-1^].

In Faizi model, photoinhibition is accounted for via the photodamage of the photosynthetic unit. The active state of photosynthetic unit (*P*^∗^) is damaged at high light intensities, then its protein is degraded to amino acid. The light inhibition in our GBA model is accounted for by an inhibition constant 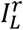 corresponding to the external light intensity *x*_L_ and the ribosome reaction *R*.

We next compare the growth rate predictions at different light intensities for both GBA and Faizi model of *Synechocystis*. **Figure 2** shows the predicted growth curves at light-limited (I), light-saturated (II), and also light-inhibited (III) for two different concentrations of external inorganic carbon. We observed that the predictions of the simpler GBA model (blue) matched well the Faizi predictions (red) at the entire range of growth rates, and by consequence also the data points (black dots) used to fit the parameters in the Faizi model. From light-limited to light-saturated regime, the GBA model predicts that the growth rate of *Synechocystis* increases to maximum amount of 0.104 h^-1^ (with doubling time of 6.66 h). Then, from light-saturated to light-inhibited growth, the growth rate starts to diminish to the amount of 0.094 h^-1^ (with doubling time of 7.36 h) as excessive light inhibits the photosystem unit, resulting in a lower reaction rate, which by consequence decreases the growth rate.

**Figure 2.**
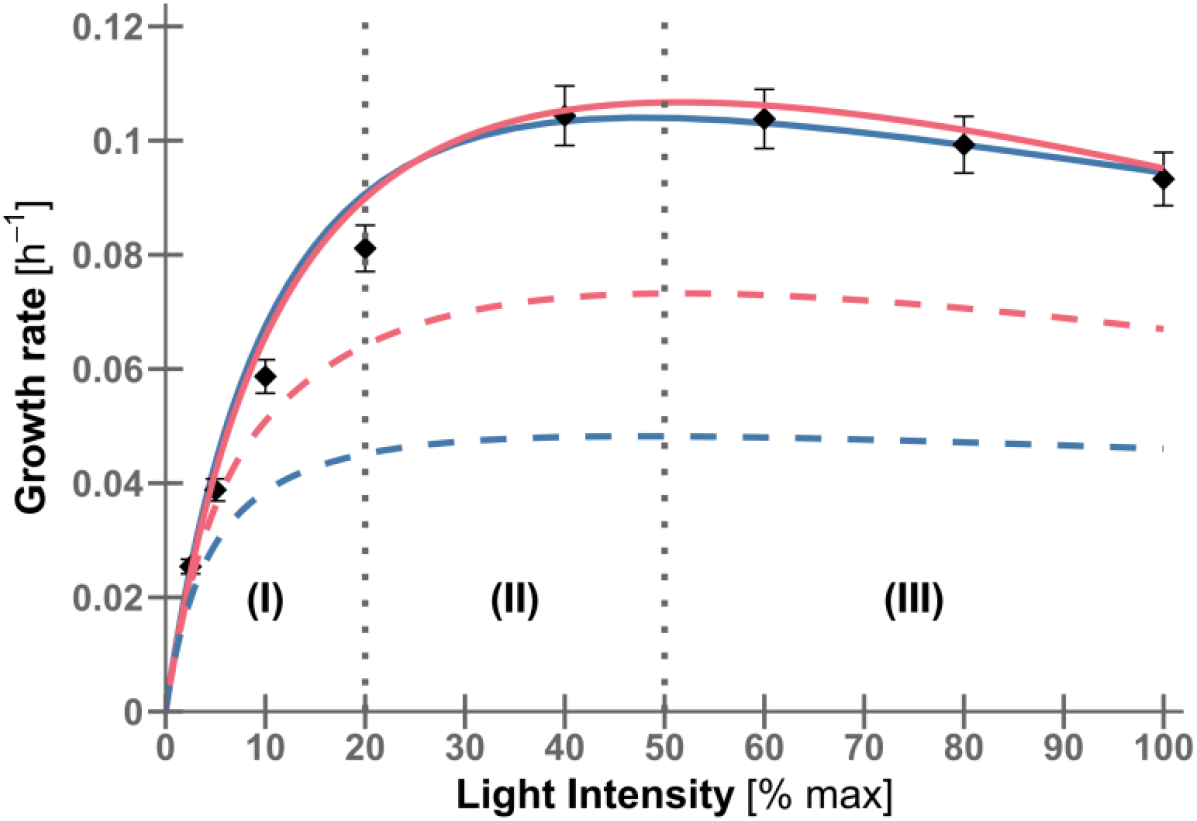
Simulated growth curve of *Synechocystis* with photoinhibition. The blue line shows the GBA growth curve for *Synechocystis* under different light intensity at high (solid line)/low (dashed line) external inorganic carbon. In comparison, the red line shows the simulation result by the Faizi model. Three different growth regimes are depicted as light-limited (I), light-saturated (II), and light-inhibited (III). The experimental steady-state growth rate for *Synechocystis* (black dots) (Faizi et al., 2018) are well explained by both models.

Next, we also compare the predicted proteome allocation for both models under different light intensities. **Figure 3** shows the predicted proteome fractions allocated to each reaction in the models. Again, the GBA predictions (blue) matched well the predictions by the Faizi model (red).

**Figure 3.**
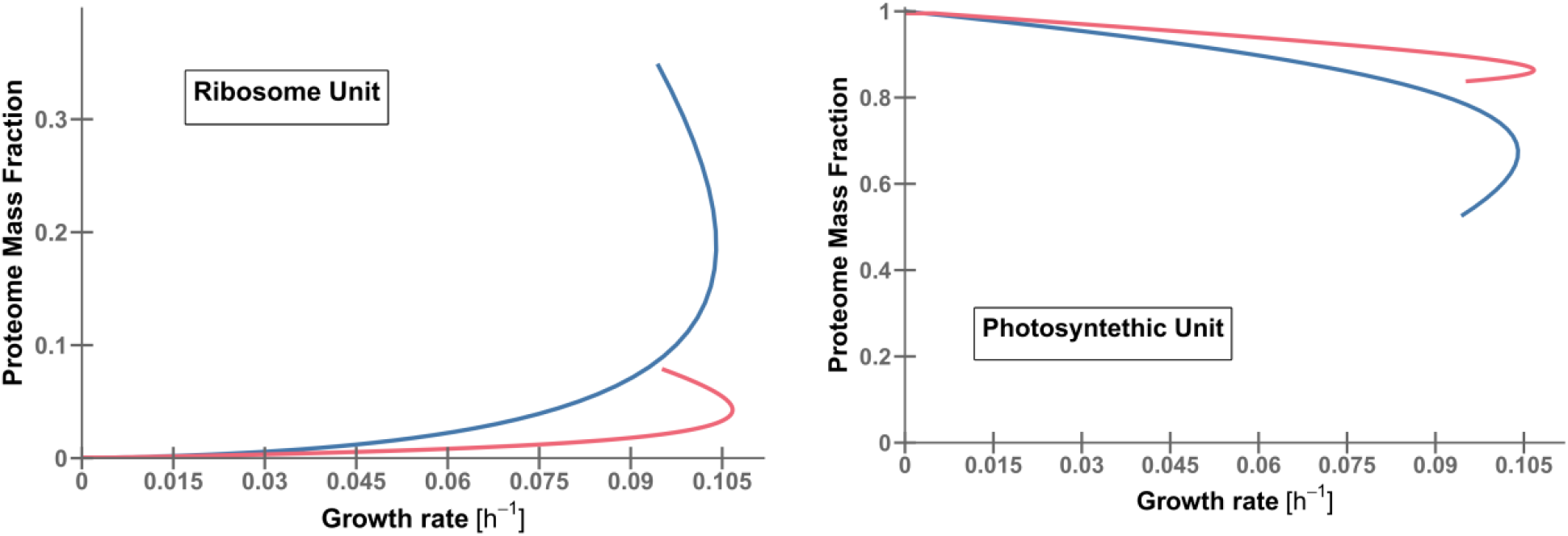
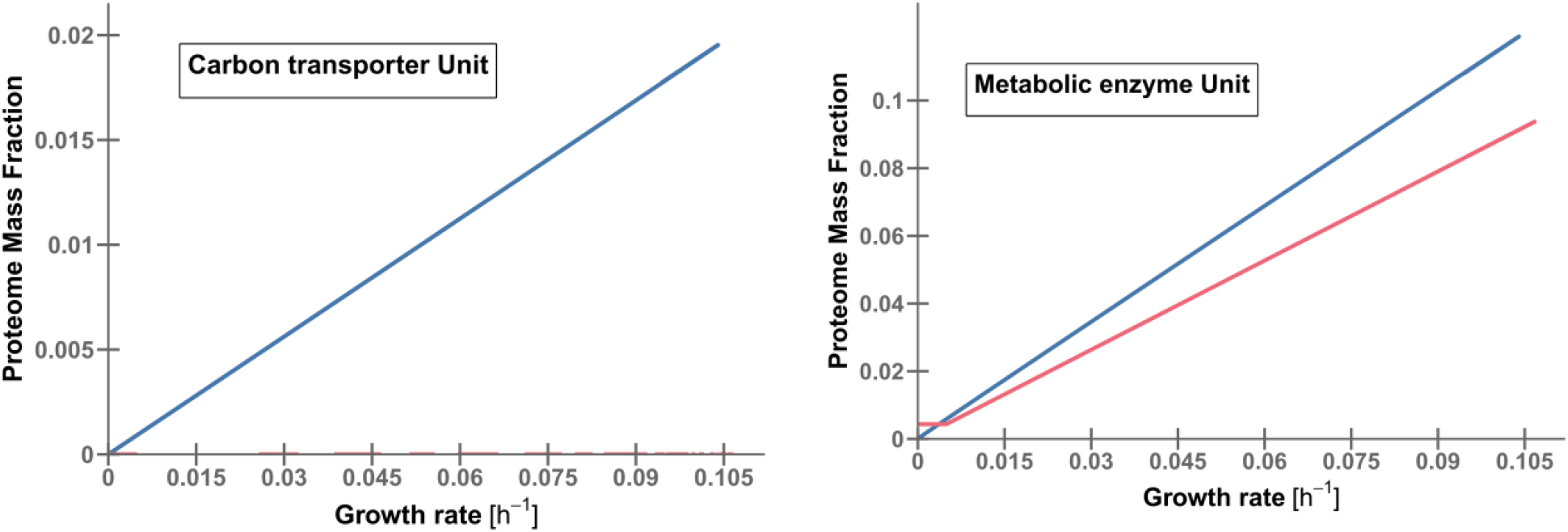
The predicted proteome allocation in the GBA model (blue) and in the Faizi model (red), including the effect of photoinhibition. The simulation data of carbon transporter for Faizi model is shown overlapping the x-axis.

**Figure 4.**
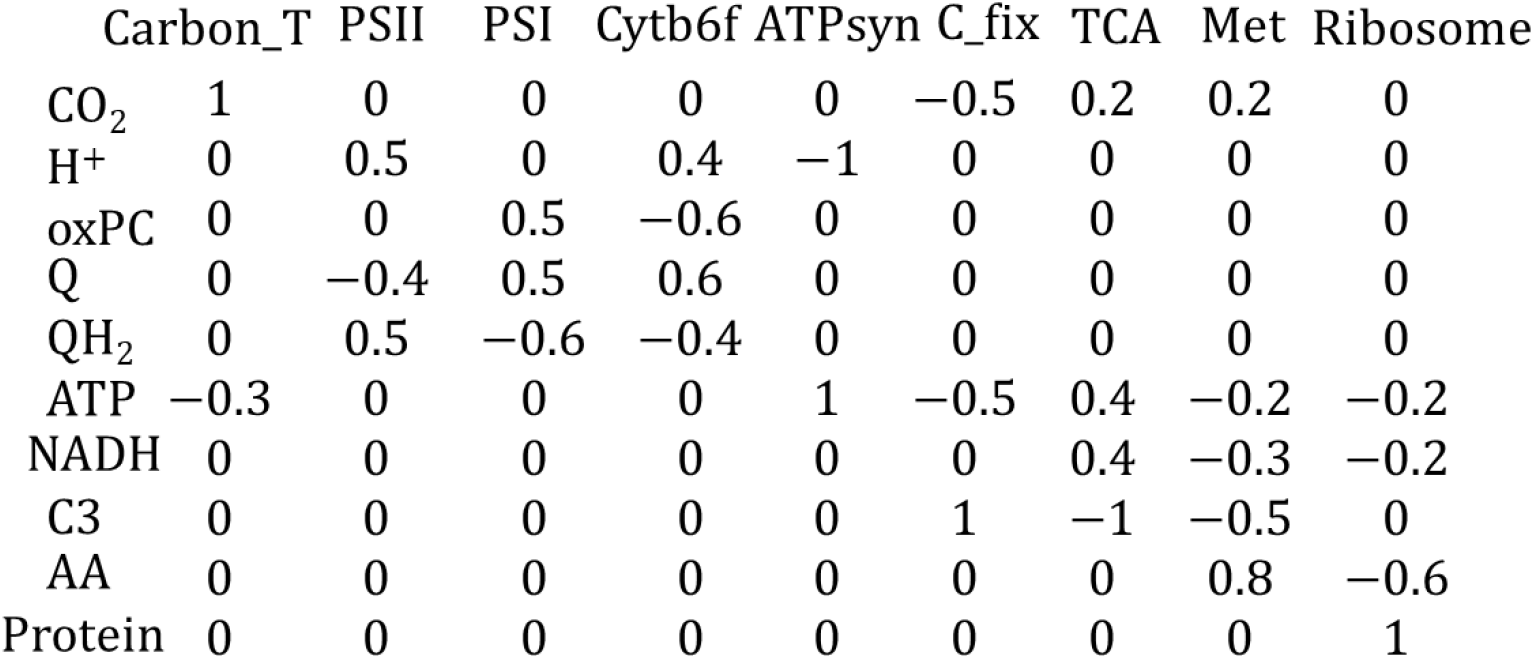
The mass fraction matrix for the second GBA model, extending the first model by including new reactions and new metabolites.

In the Faizi model, specific kinetic rate laws and mass conservation were defined for each reaction based on their function. In this type of modeling scheme, any extension of the model requires the rewriting and inclusion of new equations. In the GBA framework, the extension of any model depends only on the inclusion of new columns and rows on the matrix M, and corresponding inclusion of kinetic parameters *K*_m_, *k*_cat_ for the new reactions. This convenient strategy is explored new to build a second, more detailed GBA model of *Synechocystis*.

### Development of a second GBA model for examining photosynthesis components in cyanobacteria

Our first GBA model successfully replicated the basic phototrophic growth behavior of cyanobacteria, including the observed decrease in growth rate at high light intensities. As light intensity increases, the GBA model predicts a decrease in the proteome fraction allocated to the photosynthesis unit, and it decreases further in the photo-inhibited regime, while the growth rate decreases along with increasing in light intensity. We next examine the photosynthesis in more detail by developing a new GBA model that separates photosynthesis into four reactions: Photosystem I, Photosystem II, Cytochrome b6, and ATP Synthase. The model also includes reactions for Carbon Fixation, TCA Cycle, Metabolism, and the Ribosome. **Figure 3** presents the mass fraction matrix *M* of this new model

This second GBA model predicts growth rates at different light intensities that are almost indistinguishable from the first model predictions (not shown). This new model has however a more detailed prediction of the proteome allocation into 9 different reactions at different light intensities (see **Figure 5**), in comparison with the previous four reactions in the first model. As light intensity increases from the limited to the saturated regime, the photosynthetic mass fraction unit behaves differently from the first model. The light harvesting components, represented by the PSI and PSII proteome sectors, decrease as growth rate increases until both reach the light-saturated level. Conversely, the protein allocation to the ATPsynthase and Cytochrome b6 units increases with increasing light intensity up to this level. The ribosome proteome allocation follows an almost identical pattern as for the first model. From the light-saturated to the light-inhibited level, a downward kink was observed in the light-harvesting sectors of PSI and PSII. The PSI and PSII proteome sectors continue to decrease as growth rate decreases. Meanwhile, ATPsynthase and Cytochrome b6, which had previously increased with growth rate from the light-limited to light-saturated level, experience a downward kink as they enter the light-inhibited level. In accordance with previous observations in proteomics studies of *Synechocystis* under various growth conditions (Jahn et al., 2018; Zavřel et al., 2019), the light harvesting sectors decrease with increasing light intensity, while ATPsynthase and Cytochrome b6 increase.

**Figure 5.**
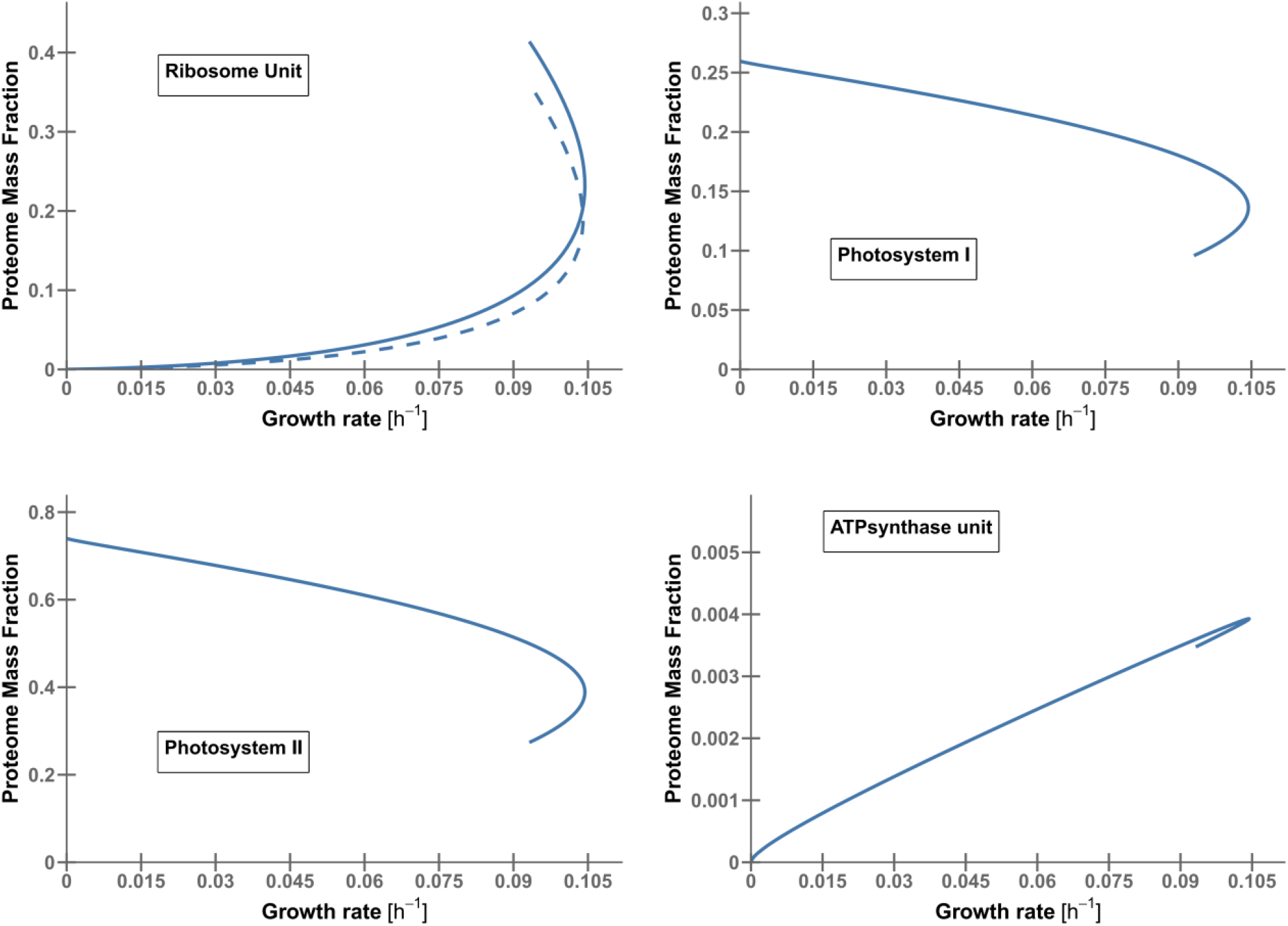
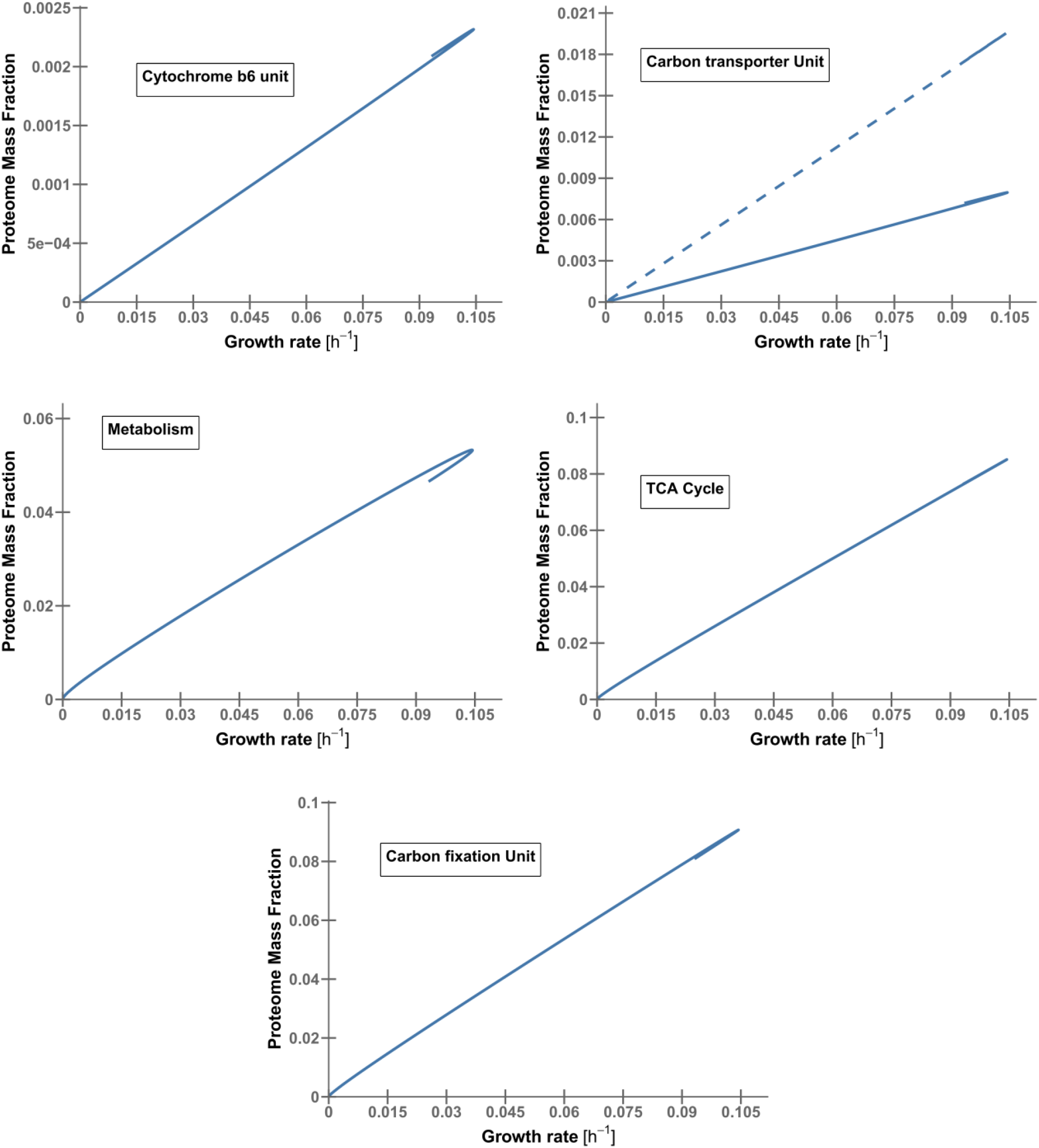
Resource allocation of *Synechocystis* in the extended model in comparison to proteomics data. The blue line (solid) shows the allocations of different proteome sectors under variations of light intensity and constant external inorganic carbon concentration. The dashed line compares the simulation results by the first GBA model with the second GBA model (solid) for the common reactions.

## Discussion

Cyanobacteria are the only prokaryotes capable of oxygenic photosynthesis, and important model systems. Most common models of cyanobacterial growth ignore nonlinear kinetic rate laws and are hence of limited use for the design of novel bioengineering strategies. Here, we have built on previous coarse-grained models for cyanobacteria (Faizi et al., 2018) and on theoretical advances in balanced growth modeling (Dourado et al., 2022) to construct simple but potentially useful models of cyanobacterial physiology, which can explain its entire resource allocation exclusively from first principles. To extend our understanding of the optimal proteome allocation in this organism, we developed a second GBA model incorporating the four key components of photosynthesis. Our GBA model demonstrated qualitative agreement with experimental findings from (Zavřel et al., 2019), whereby we observed consistent trends in proteomics data. Specifically, our model predicted a decrease in the proteome sectors of PSI and PSII, and an increase in ATP synthase and cytochrome b6 levels as the growth rate transitioned from a light-limited to a light-saturated regime, in line with the experimental results. Additionally, our model captured an upward kink in the ribosomal protein sector with decreasing growth rate, consistent with the trend observed in the experimental data(Zavřel et al., 2019).

The two GBA models presented here are the proof of concept that this type of modelling is capable of producing important results in the area of kinetic models. By providing minimal, transparent and biologically meaningful framework, we can relate the resource allocation between major protein pools and growth rate under different nutrient availability. It is constructed in such a way that the internal allocation is expressed in terms of quantifiable values. Furthermore, these quantifiable values can be grouped into categories by their functions in the cellular resource allocation models and it enables us to simply interpret the trends of interest. As seen in (Faizi et al., 2018), kinetic models are powerful tool to understand global patterns of cellular resource allocation. Although we have constructed and simulated our model around the data of Faizi for *Synechocystis*, the framework can be easily applied to other organisms as well. While other organisms need different parameter values (like different kinetic parameters or cellular density), to account for various resource allocation strategies, the fundamental framework is general and derived from basic principles. In the GBA framework, models are built from only a minimal set of parameters and a minimal set of equations defining the underlying nonlinear optimization problem.

Despite the simplicity in our assumptions, we have shown that a GBA model is capable of similar predictions as the model by (Faizi et al., 2018). This approach represents a potential advantage in constructing larger models of cyanobacteria, in which the numerical solution of the corresponding nonlinear optimization is facilitated by the simplest mathematical formulation possible. Recent studies suggested that *Synechocystis* as a model organism can introduce novel products in biotechnology and as a potential microbial cell factory(Blanc-Garin et al., 2022; Yu et al., 2013). Thus, GBA models of this organism provide a new tool to study the direct conversion of CO_2_and light to value-added chemicals and fuels, contributing to the new field of blue bioeconomy.

## Materials and methods

### Growth balance analysis framework

In growth balance analysis, the optimization problem is defined as finding the optimum growth rate (μ) subject to non-negativity constraints on metabolite and protein concentrations by varying the flux fraction ***f*** : = *v*/ μρ. Moreover, the balanced growth model at steady-state is specified by the following constraints:

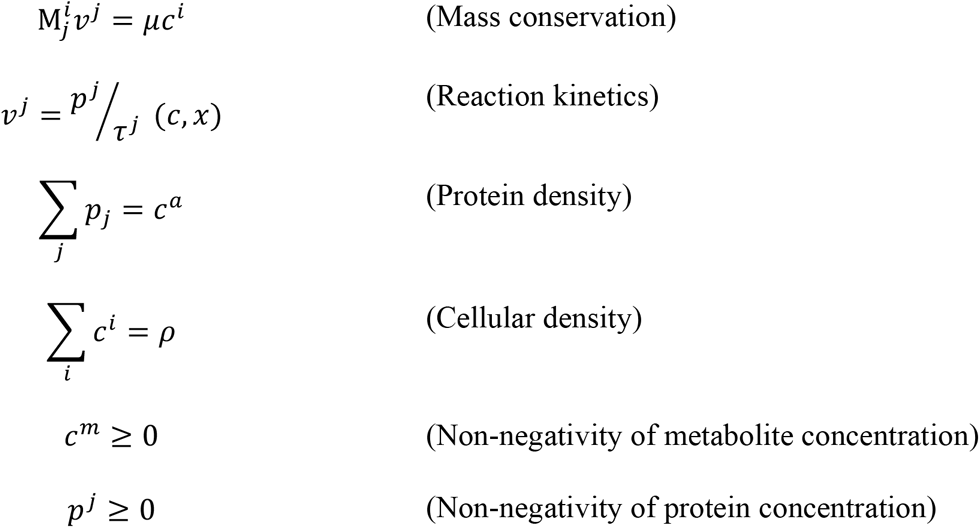

It is noteworthy to mention that the normalization of **M** results in the expression of protein concentrations (*p*_*j*_) and reactants (*c*^*i*^) in units of [g L^-1^]. Accordingly, fluxes ([g L^-1^ h^-1^]) and kinetic parameters must be represented in mass units. For instance, Michaelis constants (*K*_m_) are expressed in [g L^-1^] and turnover numbers (*k*_cat_) are represented as the amount of product per unit of protein per unit of time, resulting in units of [h^-1^]. The cell density (ρ, [g L^-1^]) is defined as the sum of all metabolite and protein concentrations, which is assumed to be constant. The comprehensive explanation of the growth balance analysis framework is provided in the original publication by (Dourado & Lercher, 2020), along with its supplementary material.

### Construction of a GBA model of *Synechocystis* sp. PCC6803 based on the model by Faizi et al

The cyanobacterial cell model presented in this study builds upon Faizi’s model in the growth balance analysis framework, but has been adapted and expanded to incorporate experimental proteomics data and better reflect the realistic characteristics of a cyanobacterial cell. The normalized mass stoichiometry of the model is defined as the stoichiometric matrix S, containing rows for reactants, is multiplied by the respective molecular mass. Then, we normalized the column so that the sum of negative entries is equal to -1 and the sum of positive entries is equal to +1 preserving mass conservation in reactions(Dourado et al., 2022). To determine the mass of protein classes in our model, we used the reference proteome of *Synechocystis* from UniProt (Bateman et al., 2017). The kinetic data (*K*_m_ and *k*_cat_values) were obtained from the BRENDA (Schomburg et al., 2013) database and then converted into [g L^-1^] for *K*_m_ and [h^-1^] for *k*_cat_. For each enzyme class, we queried the BRENDA database using the enzyme commission (EC) number of reactions in each class to find the value of the wild-type enzymes. Whenever possible, we preferred values from *Synechocystis* or other cyanobacterial cells, otherwise we directly sourced them from previously published model(Faizi et al., 2018), as indicated in the parameters section of the methods. We considered the rate-limiting *k*_cat_ for each enzyme class upon availability of various *k*_cat_ values. It is noteworthy that unlike GBA, Faizi’s model does not incorporate molecular weights in their formulation due to their molar [M] units. Therefore in our model, molecular weights were estimated in a way to fit the parameters with experimental growth curve data. Overall, the model encompasses 6 metabolites, 4 reactions, and a total of 12 parameters.

### Description of the new model

The new model is created following our implementation of cyanobacterial phototrophic growth. In simple terms, the model uses two inputs: a carbon source (e.g. CO_2_) and light, which serves as the energy source. The light is absorbed by the photosystem II and photosystem I light harvesting complexes, allowing the production of energy through ATPsynthase and the electron transport chain Cytochrome b6. The carbon source is taken in through a carbon transporter, and incorporate it into organic molecules through the process of carbon fixation, which is performed by the Calvin-Benson cycle. These are then utilized in the TCA cycle and also ribosome enzymes during protein translation. The model encompasses 12 metabolites, 9 reactions with 23 parameters, and it is noteworthy that even upon expansion, the original formulation remains unchanged.

### Model implementation

In this study, we used growth balance analysis (GBA) to simulate growth and resource allocation in cyanobacteria. R v4.1 programming language was used for implementation of balanced growth optimization problem using NLopt library. The optimization problem is solved using the augmented Lagrangian methods (AUGLAG) and Limited-memory Broyden–Fletcher–Goldfarb– Shanno (LBFGS) algorithms, which is a free and open-source software for nonlinear optimization. The models are presented in the Excel format in **Supplementary File 1**, and the R script necessary for running the simulations can be obtained from the (Dourado et al., 2022).

### Parameters of GBA models for cyanobacteria

#### 1. Stoichiometric coefficients

We started from the stoichiometric matrix (S), described by Faizi to construct our normalized mass fraction (M) in GBA model. We note that the model of Faizi does not include molecular weights of metabolites, therefore the molecular weights of metabolites should be estimated to normalize our mass fraction matrix (M). Moreover, to simplify the construction of matrix M, we estimated the entries based on the stoichiometric matrix (S) of (Faizi et al., 2018), and chose simple decimal numbers, reflecting the overall proportions of each of the columns in S.

#### 2. Molecular mass of protein classes

In order to ascertain the mass distribution of protein classes within our model (T, PSU, M, R, for the first model, and PSI, PSII, Cytb6f, ATPsynthase, carbon_transporter, carbon fixation, TCA cycle, metabolism, and Ribosome, for the second model), we employed the reference proteome of *Synechocystis* obtained from UniProt. Protein classes for the first model were obtained from the study conducted by (Zavřel et al., 2019), while protein classifications as described in (Faizi et al., 2018) along with corresponding Uniprot IDs, were utilized to expand our model to incorporate the relevant proteins. Subsequently, we systematically classified all the proteins into their respective protein categories, and then aggregated the masses of proteins within each class to derive the overall mass of each protein class. Further elaboration on the characteristics and attributes of each protein class can be found in **Supplementary Table**

#### 3. Molecular mass of metabolites

For the molecular masses of metabolites, we used 1 [g mol^-1^] for “e”, 52.5 [g mol^-1^] for “ci”(corresponding to average molecular mass of HCO_3_ and CO_2_), 36 [g mol^-1^] for “c3” (corresponding to three times of molecular mass of carbon), and 132 [g mol^-1^] for “aa” (as the average molecular mass among all amino acids(Schmidt et al., 2016)).

### 4. Kinetic parameters

#### 4.1. Carbon transporter

The turnover number (*k*_cat_) of carbon in *Synechocystis* is reported 45360 [h^-1^] in (Dornmair, Overath, & Jahnig, 1989). In order to normalize this value by mass, we multiplied by product molecular mass and then divided by the mass of enzyme, which resulted in 8.434 [h^-1^]. Besides, the Michaelis constant (*K*_m_) is also reported 15 [μM] in (Omata, Takahashi, Yamaguchi, & Nishimura, 2002), therefore we multiplied it by its substrate molecular weight, resulted in 0.0007875 [g L^-1^].

#### 4.2. Ribosome

The turnover number of ribosome for *Synechocystis* is 22 [s^-1^] (Bremer & Dennis, 2008),and the Michaelis constant for amino acids and energy is estimated as 100000 [molec cell^-1^] in (Faizi et al., 2018), which by mass normalization resulted in 13.16 [h^-1^], 0.000097812 [g L^-1^], and 0.000000741 [g L^-1^], respectively.

#### 4.3. Metabolism

The turnover number of metabolism for *Synechocystis* is reported 20 [s^-1^] in (Marcus, Altman-Gueta, Finkler, & Gurevitz, 2005), and the *K*_m_ value for internal carbon is 0.00018 [M], which results in 0.794 [h^-1^] and 0.00648 [g L^-1^]. We note that the *K*_m_= 0.000000741 [g L^-1^] for energy unit in all reactions are chosen equal for simplicity.

#### 4.4. Photosynthesis

The turnover number of photosynthetic unit is 250 [s^-1^] as reported in (Milo & Phillips, 2015), which results in 0.105 [h^-1^] and the *K*_m_ value is estimated as 0.00000741 [g L^-1^].

### 5. Second model

In this model, we followed the same procedure for calculating our model parameters. Here, we categorized the protein classes based on the new reactions in the model. Similarly, the detailed description of each protein class is provided in **Supplementary Table 2**.

#### 5.1. Mass normalized fraction

We constructed the extended model of *Synechocystis*, based on our first model to incorporate major components of photosynthesis in our model. The normalized mass fraction of can be found in the N section of our model (**Supplementary File 1**).

#### 5.2. Kinetic parameters

##### 5.2.1. Carbon transporter

The turnover number of carbon transporter (*k*_cat_) and Michaelis constant of carbon transporter is taken from the first model. The *K*_m_ value for ATP requirement to import carbon is 0.0081 [g L^-1^].

##### 5.2.2. Photosystem II

The only EC number assigned to photosystem II is 1.10.3.9. We queried the respective EC number in BRENDA to fetch the kinetic parameters for photosystem II unit. *K*_m_ value for Q (plastoquinone) is determined as 0.01236 [g L^-1^] and the maximum turnover number is 0.571 [h^-1^].

##### 5.2.3 Photosystem I

The unique EC number available for photosystem I was (1.97.1.12). The maximal turnover number for photosystem I is estimated as 0.68 [h^-1^] and the Michaelis constant for reduced plastoquinone (QH_2_) is 0.0002 [g L^-1^]. For simplicity, the *K*_m_ value for both photosystem II and I was determined as 0.000018 [g L^-1^].

##### 5.2.4. Cytochrome b6

The unique EC number available for cytochrome b6 unit was 7.1.1.6. The turnover number for this unit is 19.17 [h^-1^] and the *K*_m_ value for both oxidized plastocyanin (oxPC) and reduced plastoquinone (QH_2_) was determined as 0.000009 [g L^-1^].

##### 5.2.5. ATP synthase

The unique EC number for ATPsynthase unit was 7.1.2.2. The *K*_m_ value for proton (H^+^) in ATPsynthase unit was determined 0.0425 [g L^-1^] and *k*_cat_ value was estimated 25 [h^-1^].

##### 5.2.6. Carbon fixation

We utilized the EC. Number of 4.1.1.39 for RuBisCo as a rate limiting-step in metabolism in this protein class. Therefore, the *K*_m_ value for CO_2_ and ATP was determined 0.0088 [g L^-1^] and 0.1008 [g L^-1^], respectively. Hence, the *k*_cat_ was determined 1.783 [h^-1^].

##### 5.2.7. TCA cycle

The only EC number available for this class and for *Synechocystis* in BRENDA is 1.1.1.37. We obtained from BRENDA the corresponding *K*_m_ value for internal carbon (C3) equals to 0.145 [g L^-1^] and the *k*_cat_ equals 1.43 [h^-1^].

##### 5.2.8. Metabolism

The turnover number (*k*_cat_) and Michaelis constant of metabolism unit is taken from first model. Therefore, the *k*_cat_ is 1.6 [h^-1^] and *K*_m_ value for C3, ATP and NADH are 0.00648 [g L^-1^], 0.091 [g L^-1^], and 0.1193 [g L^-1^], respectively.

##### 5.2.9. Ribosome

The turnover number (*k*_cat_) and Michaelis constant of metabolism unit is obtained from the first model. Thus, the *k*_cat_ is 13.16 [h^-1^] and *K*_m_ value for AA, NADH, and ATP are 0.0000097812 [g L^-1^], 0.064 [g L^-1^], and 0.049 [g L^-1^], respectively.

#### 5.3. Cell densities

We obtained the cell density data from Faizi with the value of 250 [g L^-1^] and it is assumed to be constant for all the conditions.

## References

Babele, P. K., Kumar, J., & Chaturvedi, V. (2019). Proteomic De-regulation in cyanobacteria in response to abiotic stresses. Frontiers in Microbiology, 10(JUN), 1– 22. https://doi.org/10.3389/fmicb.2019.01315

Bateman, A., Martin, M. J., O’Donovan, C., Magrane, M., Alpi, E., Antunes, R., … Zhang, J. (2017). UniProt: The universal protein knowledgebase. Nucleic Acids Research, 45(D1), D158–D169. https://doi.org/10.1093/nar/gkw1099

Blanc-Garin, V., Chenebault, C., Diaz-Santos, E., Vincent, M., Sassi, J. F., Cassier-Chauvat, C., & Chauvat, F. (2022). Exploring the potential of the model cyanobacterium Synechocystis PCC 6803 for the photosynthetic production of various high-value terpenes. Biotechnology for Biofuels and Bioproducts, 15(1), 1–11. https://doi.org/10.1186/s13068-022-02211-0

Bremer, H., & Dennis, P. P. (2008). Modulation of Chemical Composition and Other Parameters of the Cell at Different Exponential Growth Rates. EcoSal Plus, 3(1). https://doi.org/10.1128/ecosal.5.2.3

Burnap, R. L. (2015). Systems and photosystems: Cellular limits of autotrophic productivity in cyanobacteria. Frontiers in Bioengineering and Biotechnology, 3(JAN), 1–13. https://doi.org/10.3389/fbioe.2015.00001

Dornmair, K., Overath, P., & Jahnig, F. (1989). Fast measurement of galactoside transport by lactose permease. Journal of Biological Chemistry, 264(1), 342–346. https://doi.org/10.1016/s0021-9258(17)31263-2

Dourado, H., & Lercher, M. J. (2020). An analytical theory of balanced cellular growth. Nature Communications, 11(1). https://doi.org/10.1038/s41467-020-14751-w

Dourado, H., Liebermeister, W., Ebenhöh, O., & Lercher, M. J. (2022). Growth Mechanics: General principles of optimal cellular resource allocation in balanced growth. BioRxiv, 2022.10.27.514082. Retrieved from https://www.biorxiv.org/content/10.1101/2022.10.27.514082v1%0A https://www.biorxiv.org/content/10.1101/2022.10.27.514082v1.abstract

Faizi, M., Zavřel, T., Loureiro, C., Červený, J., & Steuer, R. (2018). A model of optimal protein allocation during phototrophic growth. BioSystems, 166, 26–36. https://doi.org/10.1016/j.biosystems.2018.02.004

Goelzer, A., Fromion, V., & Scorletti, G. (2011). Cell design in bacteria as a convex optimization problem. Automatica, 47(6), 1210–1218. https://doi.org/10.1016/j.automatica.2011.02.038

Jahn, M., Vialas, V., Karlsen, J., Maddalo, G., Edfors, F., Forsström, B., … Hudson, E. P. (2018). Growth of Cyanobacteria Is Constrained by the Abundance of Light and Carbon Assimilation Proteins. Cell Reports, 25(2), 478–486.e8. https://doi.org/10.1016/j.celrep.2018.09.040

Liebermeister, W., & Klipp, E. (2006). Bringing metabolic networks to life: Convenience rate law and thermodynamic constraints. Theoretical Biology and Medical Modelling, 3. https://doi.org/10.1186/1742-4682-3-41

Marcus, Y., Altman-Gueta, H., Finkler, A., & Gurevitz, M. (2005). Mutagenesis at two distinct phosphate-binding sites unravels their differential roles in regulation of Rubisco activation and catalysis. Journal of Bacteriology, 187(12), 4222–4228. https://doi.org/10.1128/JB.187.12.4222-4228.2005

Matthias, K., Klähn, S., Ingeborg, S., Jasper, K. F. M., Wolfgang, R. H., & Björn, V. (2014). Comparative analysis of the primary transcriptome of synechocystis sp. PCC 6803. DNA Research, 21(5), 527–539. https://doi.org/10.1093/dnares/dsu018

Milo, R., & Phillips, R. (2015). Cell Biology by the Numbers. Cell Biology by the Numbers. https://doi.org/10.1201/9780429258770/CELL-BIOLOGY-NUMBERS-RON-MILO-ROB-PHILLIPS

Molenaar, D., Van Berlo, R., De Ridder, D., & Teusink, B. (2009). Shifts in growth strategies reflect tradeoffs in cellular economics. Molecular Systems Biology, 5(323), 1–10. https://doi.org/10.1038/msb.2009.82

O’Brien, E. J., Lerman, J. A., Chang, R. L., Hyduke, D. R., & Palsson, B. (2013). Genome-scale models of metabolism and gene expression extend and refine growth phenotype prediction. Molecular Systems Biology, 9(693). https://doi.org/10.1038/msb.2013.52

Omata, T., Takahashi, Y., Yamaguchi, O., & Nishimura, T. (2002). Structure, function and regulation of the cyanobacterial high-affinity bicarbonate transporter, BCT1. Functional Plant Biology, 29(3), 151–159. Retrieved from https://doi.org/10.1071/PP01215

Sánchez, B. J., Zhang, C., Nilsson, A., Lahtvee, P., Kerkhoven, E. J., & Nielsen, J. (2017). Improving the phenotype predictions of a yeast genome-scale metabolic model by incorporating enzymatic constraints. Molecular Systems Biology, 13(8), 935. https://doi.org/10.15252/msb.20167411

Santos-Merino, M., Yun, L., & Ducat, D. C. (2023). Cyanobacteria as cell factories for the photosynthetic production of sucrose. Frontiers in Microbiology, 14(February). https://doi.org/10.3389/fmicb.2023.1126032

Schmidt, A., Kochanowski, K., Vedelaar, S., Ahrné, E., Volkmer, B., Callipo, L., … Heinemann, aM. (2016). The quantitative and condition-dependent Escherichia coli proteome. Nature Biotechnology, 34(1), 104–110. https://doi.org/10.1038/nbt.3418

Schomburg, I., Chang, A., Placzek, S., Söhngen, C., Rother, M., Lang, M., … Schomburg, D. (2013). BRENDA in 2013: Integrated reactions, kinetic data, enzyme function data, improved disease classification: New options and contents in BRENDA. Nucleic Acids Research, 41(D1), 764–772. https://doi.org/10.1093/nar/gks1049

Wortel, M. T., Noor, E., Ferris, M., Bruggeman, F. J., & Liebermeister, W. (2018). Metabolic enzyme cost explains variable trade-offs between microbial growth rate and yield. PLoS Computational Biology, 14(2), 1–21. https://doi.org/10.1371/journal.pcbi.1006010

Yu, Y., You, L., Liu, D., Hollinshead, W., Tang, Y. J., & Zhang, F. (2013). Development of synechocystis sp. PCC 6803 as a phototrophic cell factory. Marine Drugs, 11(8), 2894– 2916. https://doi.org/10.3390/md11082894

Zavřel, T., Faizi, M., Loureiro, C., Poschmann, G., Stühler, K., Sinetova, M., … Červený, J. (2019). Quantitative insights into the cyanobacterial cell economy. ELife, 8, 1–29. https://doi.org/10.7554/eLife.42508

